# Multivalency controls the growth and dynamics of a biomolecular condensate

**DOI:** 10.1101/2025.02.01.636084

**Authors:** Julian von Hofe, Jatin Abacousnac, Mechi Chen, Moeka Sasazawa, Ida Javér Kristiansen, Soren Westrey, David G. Grier, Saumya Saurabh

**Author notes:** These authors contributed equally to this work (Julian von Hofe, Jatin Abacousnac). Contributing authors;.

## Abstract

Biomolecular condensates are essential for cellular organization and function, yet understanding how chemical and physical factors govern their formation and dynamics has been limited by a lack of non-invasive measurement techniques. Conventional microscopy methods often rely on fluorescent labeling and substrate immobilization, which can perturb the intrinsic properties of condensates. To overcome these challenges, we apply label-free, contact-free holographic video microscopy to study the behavior of a condensate-forming protein *in vitro*. This technique enables rapid, high-throughput, and precise measurements of individual condensate diameters and refractive indexes, providing unprecedented insight into size distributions and dense-phase macromolecular concentrations over time. Using this method, we investigate the kinetics of droplet growth, aging, and equilibrium dynamics in the model condensate-forming protein PopZ. By systematically varying the concentration and valence of cations, we uncover how multivalent ions influence condensate organization and dynamics, a hypothesis we further test using super-resolution microscopy. Our findings reveal that PopZ droplet growth deviates from classical models such as Smoluchowski coalescence and Ostwald ripening. Instead, we show that condensate growth is consistent with gelation at the critical overlap concentration. Holographic microscopy offers significant advantages over traditional techniques, such as differential interference contrast (DIC) microscopy, delivering reproducible measurements and capturing condensate dynamics with unparalleled precision. This work highlights the power of holographic microscopy to probe the material properties and mechanistic underpinnings of biomolecular condensates, paving the way for deeper insights into their roles in synthetic systems.

## Introduction

Biomolecular condensates are viscoelastic structures that form through phase separation of associative biological polymers [1]. They have attracted a great deal of attention in biology because of their ability to create spatial organization within cells without requiring membrane-bound structures [2, 3], in medicine for their role in physiology and disease [4], in biophysics for their potential relevance to the origin of life [5], and in chemistry for their emerging applications as self-organized catalytic centers [6, 7]. Condensates can greatly increase the efficiency of the biochemical processes they encapsulate and enable cells to regulate the associated metabolic pathways, which are crucial for signal transduction, gene expression, enzyme activity and warding of environmental stresses [8, 9].

Reconstituting condensates in minimal systems has proven to be invaluable for discovering their physicochemical properties and the biological phenotypes that are associated with them [3, 10, 11]. Condensates form by partitioning macromolecules such as proteins and nucleic acids into droplets of high macromolecular concentration dispersed within a dilute phase. The concentration of macromolecules in the dense phase determines the droplets’ viscoelastic and interfacial properties and so influences their morphology [12, 13]. The dense-phase concentration also determines the biological role of the condensate by establishing the droplets’ ability to localize other macromolecules under conditions that support their biochemical activity. The size distribution of condensate droplets offers complementary insights into the mechanisms by which the condensed phase nucleates and grows within a macromolecular solution, and therefore offers insights into the tunability of the droplets’ functionality [14].

Imaging-based studies of condensate concentration, size distribution and morphology typically rely on fluorescent labeling and often require droplets to be immobilized on glass substrates. Dyes and substrates, however, can perturb the phase behavior of macromolecular solutions as well as the kinetics of droplet condensation. Label-free methods such as quantitative phase imaging [15] can be used to estimate macro-molecular concentration [16], but still require droplets to be immobilized on surfaces. Adhesion to a substrate alters the droplets’ apparent size distribution both by distorting individual droplets and also by mediating droplet fusion [17]. Scanning a surface to build statistics also limits throughput.

We overcome the limitations of conventional analytical techniques by using holographic video microscopy [18, 19] to analyze large populations of unlabeled biomolecular condensates *in vitro*. Each hologram of a micrometer-scale droplet encodes a wealth of information, including the droplet’s size, shape and refractive index. This information can be retrieved rapidly and with great precision by fitting the droplet’s hologram to a generative model based on the Lorenz-Mie theory of light scattering [18, 20]. Large populations of droplets and particles can be characterized by streaming fluid dispersions through the observation volume of a holographic microscope in a microfluidic channel [19] and analyzing their holograms in real time [21]. Holographic characterization yields precise estimates for the size and refractive index of each particle in a precisely specified measurement volume [22], and therefore yields exceptionally accurate estimates for the distributions of those quantities [23]. A particle’s refractive index, in turn, can be interpreted with effective-medium theory to obtain quantitative insights into its composition [24, 25]. Holographic characterization is free from perturbations due to labeling or substrate interactions and can amass data on thousands of particles in a matter of minutes.

Here, we apply holographic microscopy to study the assembly and growth mechanisms of PopZ, a model condensate-forming protein that regulates the cell cycle in the aquatic and soil bacterium, *Caulobacter crescentus* [3, 26, 27]. We use holographically-measured refractive indexes to infer the concentration of PopZ within individual condensate droplets and thereby to determine how the dense-phase concentration depends on the concentration and valence of the cations used to trigger phase separation. The precision and speed afforded by digital holography enable us to monitor the kinetics of condensate formation, growth and aging over time. These measurements reveal that the mean droplet diameter grows more slowly than expected for Ostwald ripening, the variance in diameter increases faster than predicted by Smoluchowski kinetics, and at late times, the growth rate aligns with self-regulated aggregation kinetics. Critically, our data suggested that PopZ condensate composition and dynamics are controlled by the multivalency of cations used, a hypothesis that we tested via super-resolution microscopy and molecular dynamics simulations. In casting new light on macromolecular condensation in a powerful model system, our results also highlight the value of holographic video microscopy for high-throughput and perturbation-free measurements of biomolecular condensates.

## Results

### Precise, substrate-free characterization of condensate properties

PopZ is an intrinsically disordered protein localized at the poles in *Caulobacter crescentus* that plays a crucial role in localizing and spatially organizing multiple regulatory proteins from the cytoplasm. The structure of PopZ includes an amphipathic *α*-helix near the N-terminal, which mediates interactions with its “guest” proteins [28]. PopZ 3 also contains an intrinsically disordered region (IDR) enriched with acidic residues (Fig. 1a) that has been suggested to control condensate fluidity [29], and C-terminal helical domains crucial for higher-order oligomerization [28, 30]. PopZ undergoes phase separation *in vivo* [27, 29, 31] where it forms condensates that exclude ribosomes and are essential for proper cell division and fitness. Owing to its highly negatively charged IDR, PopZ can be induced to form condensate droplets *in vitro* in the presence of cations, where it constitutes a versatile model system for studying macro-molecular condensation [3, 29]. Figure 1a schematically illustrates the protocol for inducing liquid-liquid phase separation in PopZ solutions by adding magnesium chloride (abbreviated as Mg^2+^, hereafter for simplicity). This produced a population of micrometer-scale condensate droplets observable with differential interference contrast (DIC) microscopy (Fig. 1a, S1a).

**Fig. 1.**
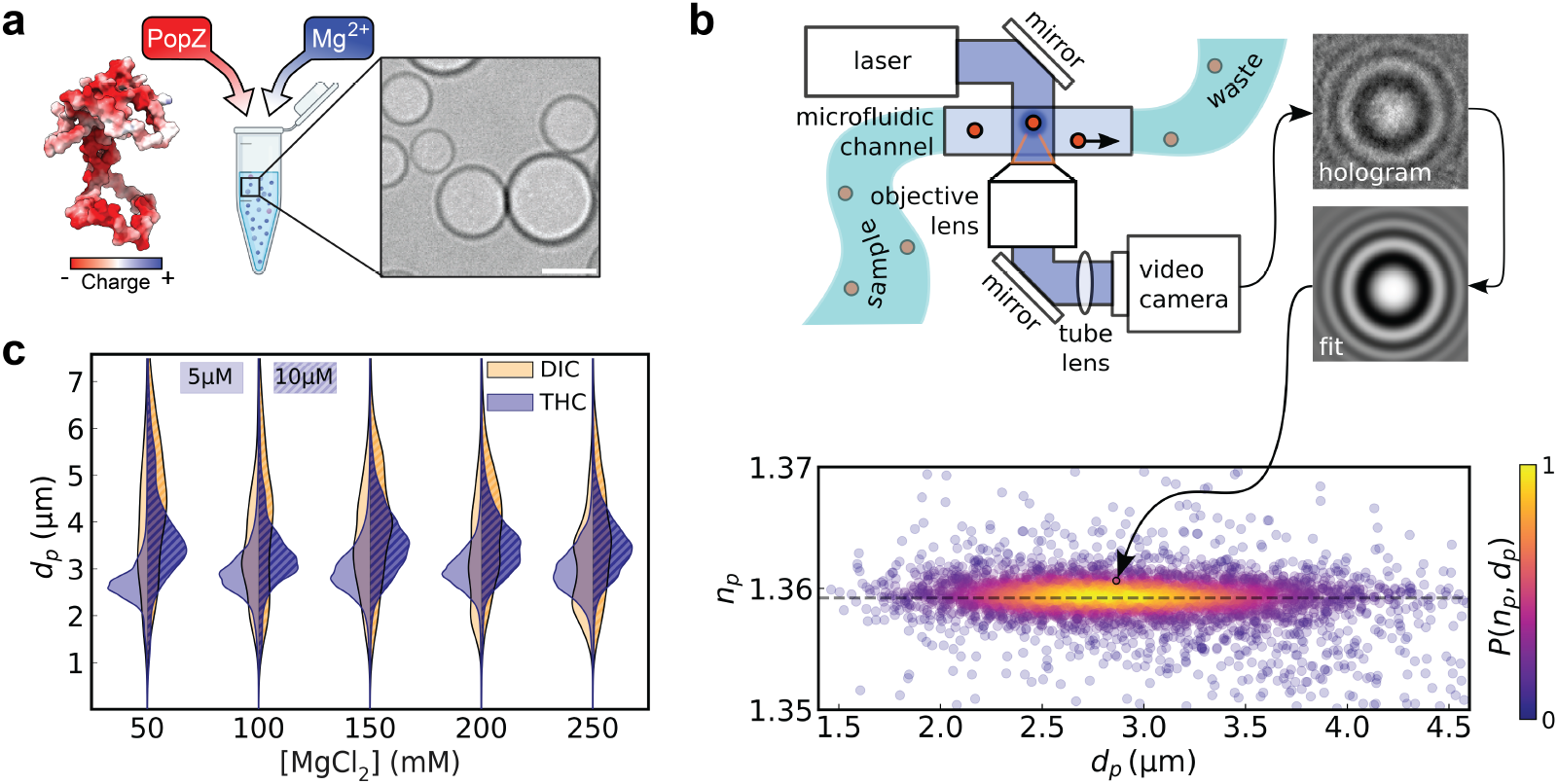
Characterization of PopZ condensates using holographic microscopy. (a) Schematic of PopZ protein condensate formation in the presence of magnesium ions (Mg^2+^), shown alongside a DIC microscopy image of PopZ condensates. The image highlights their spherical morphology. Scale bar: 5 µm. (b) Diagram of the holographic characterization setup for condensate analysis, which employs a microfluidic channel to flow samples through a laser-based holographic microscope. The objective lens captures the scattered light to produce holograms that are subsequently analyzed to extract physical parameters, including diameter *d*_*p*_, and refractive index, *n*_*p*_. The scatter plot illustrates the measured *n*_*p*_ and *d*_*p*_ values for 4383 condensate droplets, with colors representing the probability density, *P* (*n*_*p*_, *d*_*p*_). The horizontal dashed line indicates the mean refractive index, *n*_*p*_ = 1.359 21 ± 0.000 03. (c) Violin plots of PopZ condensate size distributions across a range of Mg^2+^ concentrations (50 µM to 250 µM) at two initial PopZ concentrations (50 µM and 10 µM). Holographic microscopy provides consistent size measurements without substrate effects, outperforming traditional DIC microscopy.

To analyze PopZ condensates via holographic microscopy, 30 µL of the phase-separated sample was transferred to a microfluidic channel as illustrated schematically in Fig. 1b, and the population of droplets passing through the channel in a pressure-driven flow was analyzed. The flow transports particles through the microscope’s laser beam, ensuring that each particle has an opportunity to scatter a proportion of the light. The scattered light interferes with the remainder of the laser beam, and the intensity of the magnified interference pattern constitutes a hologram of the particle that is recorded with a video camera. Provided the number density of condensates does not exceed 10^7^ mL^−1^, holograms of individual droplets are well separated in the field of view and can be analyzed with a generative model [18, 20] based on the Lorenz-Mie theory of light scattering [32] (Fig. 1b) to obtain estimates for the droplet’s diameter, *d*_*p*_, and refractive index, *n*_*p*_.

Figure 1c presents holographic characterization results from 4383 PopZ condensates, with the diameter and refractive index of each droplet being represented by a single point. Points are colored by the relative density of observations, *P* (*n*_*p*_, *d*_*p*_). The droplets in this sample have a broad distribution of diameters, ranging from *d*_*p*_ = 1 µm to 5 µm. By contrast, they have a remarkably narrow distribution of refractive indexes, *n*_*p*_ = 1.359 21 ± 0.000 03, which is noteworthy because the refractive index serves as a proxy for the concentration of protein in each droplet [24, 25, 33]. Holograms were measured with laser light at a vacuum wavelength of *λ* = 450 nm, for which the refractive index of the aqueous buffer medium is *n*_*m*_ = 1.340 ± 0.001 at room temperature. The droplets’ refractive index therefore is clearly resolved. For such samples, holographic microscopy has been shown [25] to have a detection efficiency close to unity for particles ranging in size from *d*_*p*_ = 0.5 µm to *d*_*p*_ = 10 µm. The distribution plotted in Fig. 1c therefore provides a comprehensive view of the population of droplets in the sample. The data for this measurement were acquired in the initial 2 min of an 8 min measurement window and therefore represent a snapshot of the sample’s properties. Sequences of such measurements therefore can be used to track the time evolution of the condensate droplets’ properties.

The narrowness of the refractive-index distribution and the lack of correlation between refractive index and size suggests that PopZ is distributed uniformly within each droplet and that the droplets all have the same concentration of PopZ, independent of size. This differs markedly from previously reported holographic characterization results for protein aggregates, whose inhomogeneous fractal structure yields a broad distribution of refractive indexes and a characteristic dependence of refractive index on cluster size [25, 33, 34]. Neither the droplets’ sizes nor their refractive indexes are correlated with their height in the microfluidic channel (Fig. S1b), furthermore, indicating that hydrodynamic shear has no effect on PopZ condensates’ size and shape under these conditions [21].

Whereas holographic microscopy can characterize free-flowing condensates, standard techniques such as DIC microscopy require droplets to be immobilized on substrates, which can affect their measured properties (Fig. S1c). We quantified the reproducibility of size measurements between DIC and holographic techniques by measuring the Jensen-Shannon divergence (JSD) [35] between measurement runs (Fig. S1d). Holographic size measurements yielded a JSD score of 0.02 ± 0.01, which is an order of magnitude better than the score of 0.40 ± 0.04 obtained for DIC measurements. Size distributions obtained with holographic microscopy therefore are significantly more reproducible, run-to-run. Droplet-to-droplet and run-to-run variations in DIC measurements may reflect the influence of adhesion to the immobilizing substrate on condensate phase behavior.

### Measurement of PopZ concentration in the condensed phase

Macromolecular condensation is an example of liquid-liquid phase separation mediated by intermolecular interactions [36]. Because PopZ has an abundance of acidic residues, the phase behavior of its solutions and the properties of its condensates should depend on the concentration and valence of the cations in the buffer. Accordingly, a phase diagram of PopZ in the presence of a range of Mg^2+^ concentrations revealed the presence of a phase boundary for cation concentrations greater than 50 mM (Fig. S1a). Next, we used DIC, holographic characterization, and confocal fluorescence microscopy to monitor how the physical properties of PopZ condensates depend on Mg^2+^ concentrations in the range from 50 mM to 250 mM. DIC microscopy reveals the presence of condensates at each concentration (Fig. 2a), although no differences in condensate protein concentrations can be discerned from the relative intensities of the recorded images. Fluorescent labeling is commonly used to estimate protein concentrations in the dense phase of biomolecular condensates. Accordingly, we used NHS ester conjugates of BODIPY-FL and JF646 dyes and analyzed the fluorescence intensities inside (*F*_in_) and outside (*F*_out_) condensates using confocal microscopy (Fig. S2a-c). The results reveal significant dye-dependent differences, with JF646 showing higher fluorescence intensity in the dense phase compared to BODIPY-FL, even under identical labeling conditions. While the fluorescent signal from BODIPY-FL labeled PopZ increases both inside and outside the condensates, JF646 conjugated PopZ does not show significant differences in the dense phase concentration of PopZ. For the BODIPY-FL conjugated PopZ, *F*_in_*/F*_out_ values increase with Mg^2+^ concentration, suggesting that higher ionic strength promotes labeled-protein sequestration in the dense phase (Fig. S2d). However, inconsistencies in *F*_out_ between the dyes and across conditions complicate the interpretation, highlighting how the dyes’ chemical properties influence protein partitioning.

**Fig. 2.**
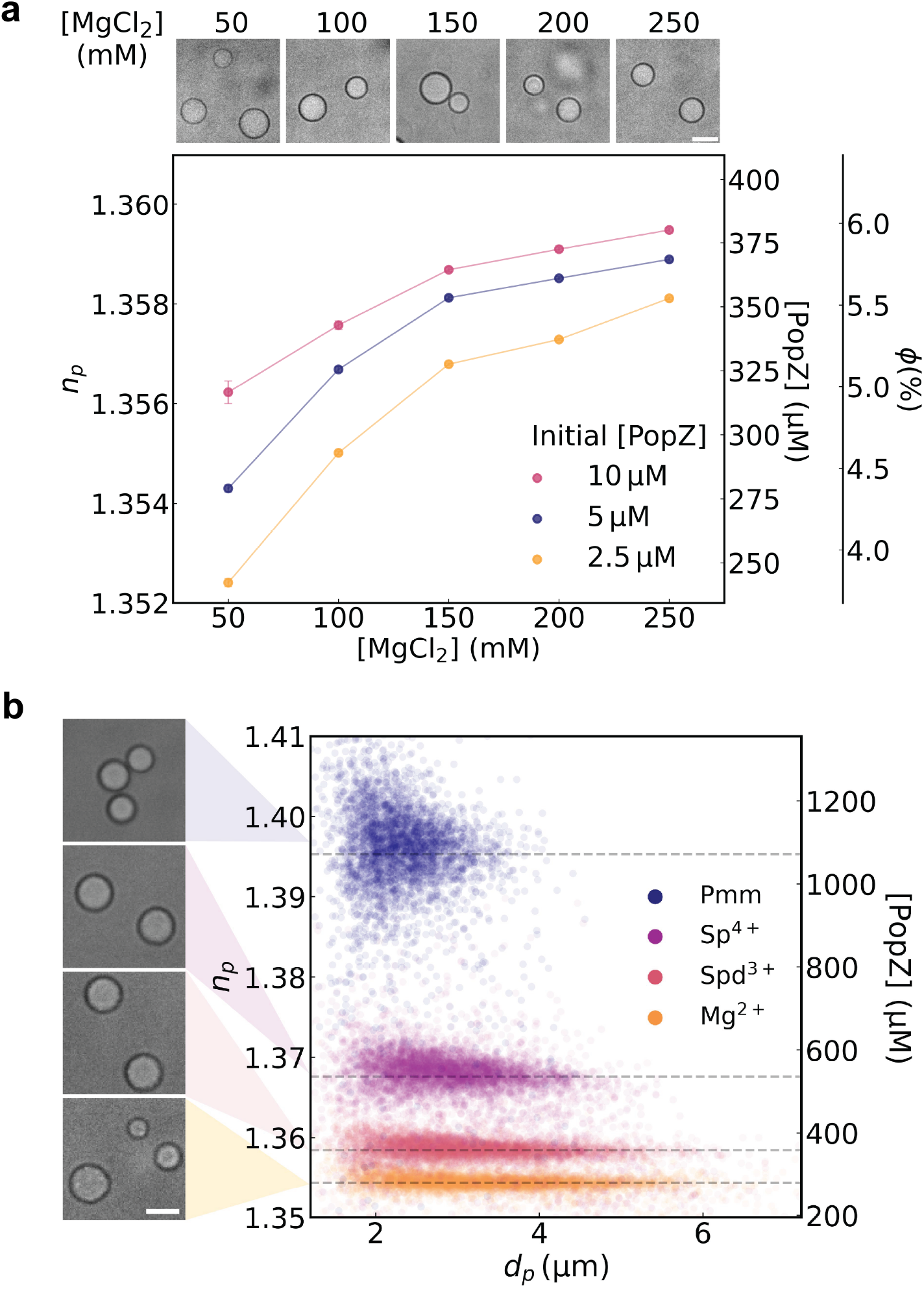
Effect of multivalent ions on the refractive index and size of PopZ condensates. (a) Top: DIC microscopy images of PopZ condensates formed at different Mg^2+^ concentrations (50 mM, 100 mM, 150 mM, 200 mM, and 250 mM). Scale bar: 5 µm. Bottom: Dependence of droplet refractive index (*n*_*p*_) on Mg^2+^ concentration for different PopZ concentrations (2.5 µM, 5 µM, and 10 µM). The secondary axes map the refractive index values onto the dense phase concentration and volume fraction, respectively, using effective medium theory according to Eqs. (10) and (12). (b) Left: DIC microscopy images of PopZ condensates formed by (from top to bottom) Pmm (50 µM), Sp^4+^ (25 mM), Spd^3+^ (33.3 mM), or Mg^2+^ (50 mM). Scale bar: 3 µm. Right: Scatter plot of condensate droplets’ diameters (*d*_*p*_) and refractive indexes (*n*_*p*_) colored by the polycation used to trigger condensation. The secondary axis maps the refractive index values onto the dense-phase concentration, using effective medium theory. Horizontal dashed lines correspond to the mean refractive index values for each ion type.

Whereas DIC and confocal microscopy offer indirect insights into condensate composition, holographic characterization directly measures the refractive indexes of individual condensate droplets. Section 5 explains how the refractive index can be used to compute the volume fraction of protein in a droplet and therefore the absolute concentration of protein in the dense phase. The basis for this precise high-throughput label-free measurement technique is illustrated in Fig. S3a. Each point in Fig. 2a represents the mean refractive index from a thousand-droplet distribution such as the example in Fig. 1c and is reproduced in triplicate. Error bars denote the standard error in the mean for the three samples. The data in Fig. 2a reveal that the concentration of PopZ in the dense phase increases both with increasing concentration of PopZ in the starting solution and also with increasing concentration of Mg^2+^. The dense-phase concentration appears to approach a plateau at higher Mg^2+^ concentrations, suggesting that the divalent ion and PopZ exhibit saturable binding. The plateau concentration appears to depend comparatively weakly on the overall concentration of PopZ in the system. The influence of Mg^2+^ concentration on the measured protein concentration in the dense phase suggests a role for cation valence in modulating the composition of the condensed phase.

We hypothesized that increasing the valence of the cation used to trigger phase separation would further increase the dense phase concentration. To test this idea, we form PopZ condensates in the presence of polyvalent cations with different valence: trivalent spermidine trichloride (Spd^3+^), tetravalent spermine tetrachloride (Sp^4+^), and a polyamidoamine dendrimer called PAMAM (Pmm) that has 16 terminal amines [37, 38]. PopZ condensates are formed at physiological pH 7.0. Under these conditions, the polyamines Spd^3+^ and Sp^4+^ are fully protonated and thus exhibit cationic valences of +3 and +4, respectively [39]. Although the degree of protonation of Pmm at pH 7.0 is not precisely known [38], this dendrimer should have an effective valence greater than that of Sp^4+^, and thus serves as an extreme case of cationic valence. To facilitate comparison, Mg^2+^, Spd^3+^, and Sp^4+^ are added at concentrations such that they result in an equivalent total positive charge. The DIC images in Fig. 2b illustrate that all four types of cations induce liquid-liquid phase separation and produce spherical condensate droplets. The data in Fig. 2b reveal that droplets are produced in roughly the same numbers with all four types of cations but that their protein content varies dramatically with cation valence.

From divalent Mg^2+^ to polyvalent Pmm, PopZ condensates undergo a four-fold increase in their concentration. To place this in context, the volume fraction of PopZ in the dense phase increases from *ϕ* = 0.04 in the presence of Mg^2+^ to *ϕ* = 0.16 in the presence of Pmm (Fig. S3b). The measured volume fraction of protein in PopZ condensates is substantially lower than the volume fraction of polymers in typical latex colloids [40], suggesting that PopZ condensates are likely to be much softer [41]. For reference, the phase diagram of PopZ as a function of each polyamine’s concentration is provided in Fig. S4. Finally, the distribution of refractive index values also broadens with higher cation valence, suggesting that polyvalent cations create more opportunities for structural heterogeneity within each droplet, a hypothesis we test later. The increase in dense-phase concentration is associated with a decrease in the average droplet diameter and an increase in the dispersity in size. When combined with the observation that droplets form at roughly the same number density in all four systems, this observation suggests that increasing cation valence does not substantially affect the rate of droplet nucleation, but subsequently fosters growth of smaller denser condensate droplets. Accordingly, we investigated next the growth mechanisms of PopZ condensates through time-resolved holographic imaging.

### Measured condensate size distributions constrain models for droplet growth

The ability of cells to control the size of condensate droplets has been proposed as a mechanism to modulate the macromolecular concentration within the droplets and thereby to regulate the droplets’ functionality [8]. Monitoring the size of condensate droplets can also be useful for diagnosing diseased states. For example, dysregulation of the sizes of nucleolar condensates has been correlated with cancer prognosis severity [42]. A recent study proposes that condensate sizes in biological systems follow either an exponential distribution or a power-law distribution depending on the relative rates of nucleation and coalescence [14]. These findings, consistent across native cellular condensates, synthetic optogenetic systems, and computer simulations, suggest a biophysical principle governing condensate formation and size regulation in cells.

To gain insight into the growth mechanisms of PopZ condensates, we use the capabilities of holographic microscopy to monitor the size distribution of condensate droplets as a function of time after phase separation is triggered by the addition of Mg^2+^. Typical results for the time-dependent size distribution, *P* (*d*_*p*_, *t*), are plotted in Fig. 3a. Quantitative analysis contraindicates droplet coalescence and Ostwald ripening as viable explanations for the observed time evolution of *P* (*d*_*p*_, *t*), and instead suggests that these droplets grow by self-regulating coagulation of macroions at the critical gel point.

**Fig. 3.**
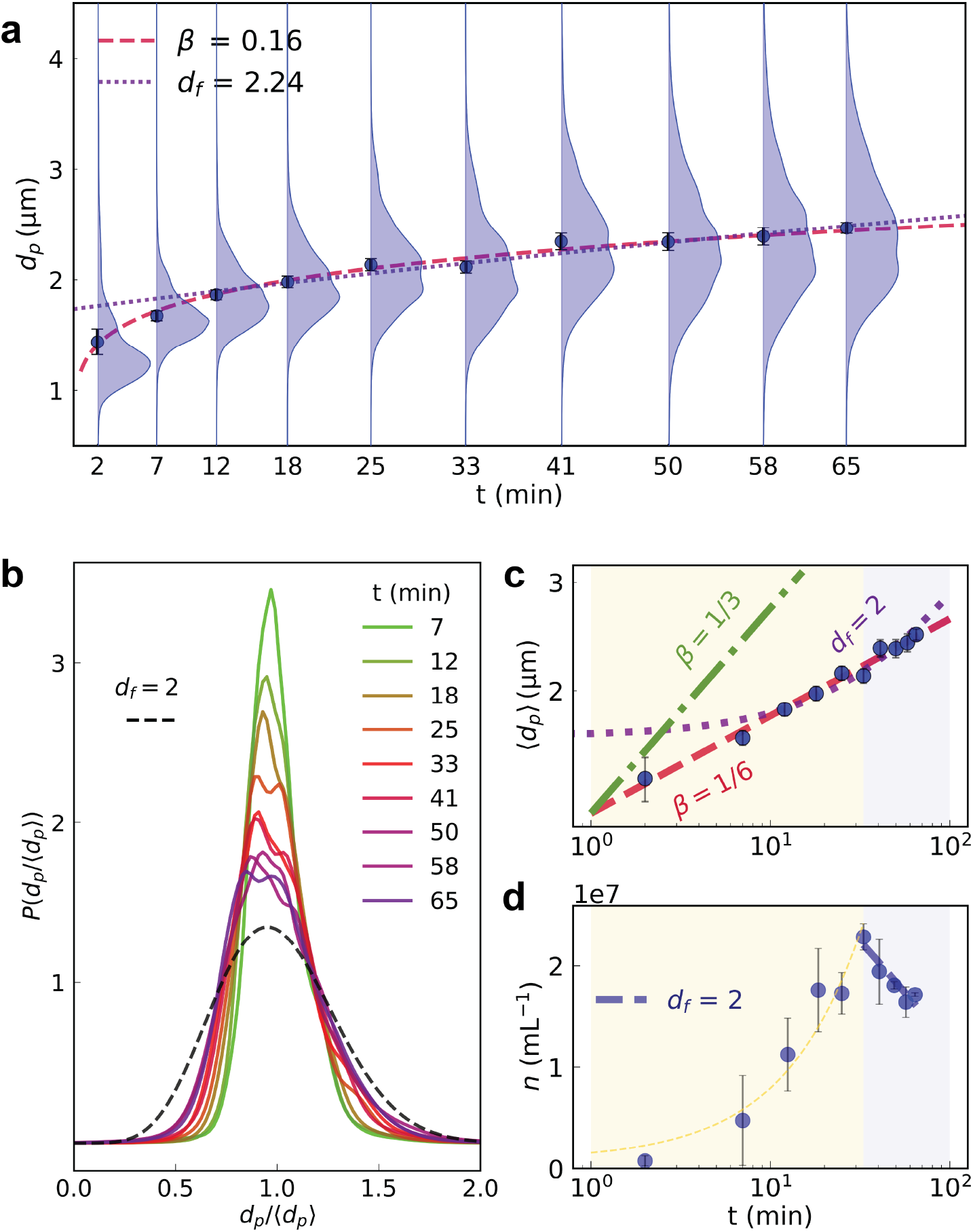
Time-dependent evolution of PopZ condensates. (a) Violin plots showing condensate size distributions, *P* (*d*_*p*_, *t*), for 2.5 µM PopZ at 50 mM Mg^2+^, at various time points, demonstrating the progressive increase in condensate size distribution over time. Discrete (dark blue) points represent the mean condensate diameter, ⟨*d*_*p*_(*t*)⟩, at each sampling time, with error bars indicating the standard error of the mean. at each time. The red dashed curve is a fit to a power law, ⟨*d*_*p*_(*t*)⟩ ∼ *t*^*β*^. (b) Probability densities, *P* (*d*_*p*_*/* ⟨*d*_*p*_⟩), of droplet diameters scaled by the mean droplet diameter at each sampling time, *t*. These distribution functions should collapse onto a single curve for a system displaying dynamic scaling characteristic of growth by droplet coalescence. (c) Log-log plot of the mean droplet diameter showing power-law scaling. The fit exponent, *β* = 0.16 ± 0.01, is inconsistent with the value of 1*/*3 expected for Ostwald ripening (green dot-dashed line). At late times, the evolution of mean droplet diameter aligns with Eq. (3) consistent with self-regulated kinetics (violet, dotted curve). (d) Droplet concentration over time, following predictions from self-regulated kinetics at late times. The initial increase in concentration suggests nucleation-driven processes.

The mean droplet diameter at each measured time interval, ⟨*d*_*p*_(*t*)⟩, is denoted by discrete points in Fig. 3a, with error bars representing the standard error in the mean. The dashed curve passing through these points suggests that droplets grow as a power law in time,

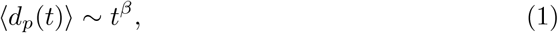

with the exponent *β* = 0.16 ± 0.01. Power-law growth is consistent with coarsening by coalescence, which is described by the Smoluchowski kinetic model [43, 44]. Assuming that droplets collide and merge at a rate that is limited by diffusion, the Smoluchowski equation predicts that their size distribution should have the form [45]

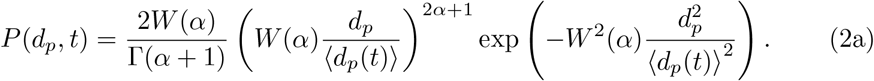

The width of this distribution,

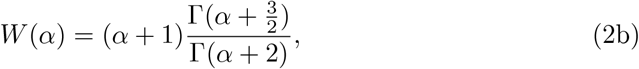

is set by the size dependence of the droplets’ diffusion coefficient,

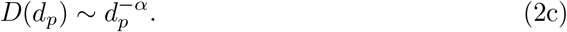

In the dynamic scaling hypothesis, the exponent *α* establishes the droplets’ growth rate:

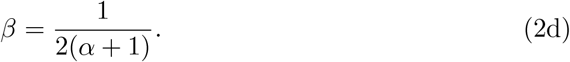

The observed value of *β* suggests *α* = 1.94 ± 0.10 which differs significantly from the standard Stokes result, *α* = 1, for solid spheres, but could be consistent with the scaling predicted for a fractal polymer gel [46].

Even if coalescence can account for the growth in the mean droplet diameter, it does not explain the observed broadening of the size distribution. Equation (2) suggests that *P* (*d*_*p*_, *t*) should collapse onto a single master curve if *d*_*p*_ is normalized by ⟨*d*_*p*_(*t*)⟩. Instead, the scaled distribution functions plotted in Fig. 3b broaden with time, and only appears to reach an asymptotic form after 45 min. This broadening is apparent because holographic microscopy builds a statistical sample rapidly enough to resolve it. The asymptotic distribution width corresponds to *α* = 3.8 ± 0.1, which differs from the dynamic scaling hypothesis and does not have an obvious interpretation in terms of the droplets’ microstructure or dynamics. This disagreement effectively rules out droplet coalescence as the dominant coarsening mechanism.

Ostwald ripening also causes coarsening with power-law time dependence [47]. In this case, however, the Lifshitz-Slyozov-Wagner theory predicts the growth exponent to be *β* = 1*/*3, which is twice the observed value. The difference is clearly resolved in Fig. 3c. Droplet growth therefore does not appear to be driven by simple diffusive exchange of macromolecules between liquid-like droplets. This is consistent with previous studies of PopZ condensates using fluorescence recovery after photobleaching *in vitro* [3] and lower mobility fractions *in vivo* [29]. The apparent absence of Ostwald ripening in this system suggests that the surface tension of PopZ condensates must be low, which also appears to be the case for condensates of coiled-coil proteins [48].

Holographic microscopy provides accurate values for the number density of condensates, *n*(*t*), throughout the accessible size range [23] and therefore tracks changes in droplet number due to processes such as nucleation and coalescence independent of changes in their size distribution. The data in Fig. 3d show that the number density of detectable condensates increases for the first half hour after condensation is triggered, suggesting that nucleation continues for several minutes after condensation is triggered. The number density declines after half an hour, which suggests that coarsening outstrips the nucleation rate thereafter.

The observed growth exponent, *β* ∼ 1*/*6 is reminiscent of the slow coagulation of macroions in the presence of multivalent counterions [49], which is a model system for the coagulation of protein chains in the presence of polyvalent cations. As condensate aggregates grow, the Coulomb barrier between them increases, reducing the likelihood of further coagulation. This “self-regulated” growth mechanism, which begins at time *t*_0_ *>* 0, predicts a logarithmic growth law for the mean droplet diameter

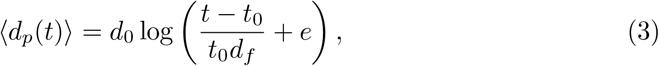

where *d*_0_ represents the mean droplet size at time *t*_0_. The model further suggests that the concentration of typical aggregates decreases logarithmically with time, following

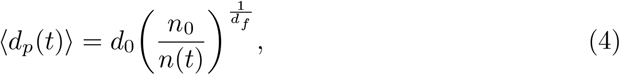

where *n*_0_ is the concentration of droplets at time *t*_0_, and *n*(*t*) is the time-dependent concentration. To test this model, we measured the droplet concentration over time, and identified two distinct regimes (Fig. 3d). At early times, the concentration increases monotonically, consistent with nucleation driven growth. Around the 30 min mark, the concentration begins to decrease steadily. In this later regime, we fit the concentration data to Eq. (4) assuming a fractal dimension *d*_*f*_ = 2. Furthermore, the logarithmic dependence of the mean droplet size, as shown in Fig. 3c, is consistent with this fractal dimension. Together, these observations suggest that at late times, the slow growth of condensates follows the predictions of self-regulated kinetics, rather than Ostwald ripening or Smoluchowski coagulation. The fractal dimension of the condensates in this regime matches that of a crosslinking polymer at the gel point, reinforcing the idea of a self-regulated growth model.

### Physicochemical perturbations drive condensates out of equilibrium

Reconstituted condensates are usually studied at or near equilibrium. Living systems, by contrast, rely on condensates’ properties far from equilibrium. To bridge this gap, we combine DIC and holographic microscopy to study how condensates respond to changes in temperature and the chemical environment. PopZ condensates imaged via DIC microscopy showed dissolution at higher temperatures (Fig. 4a), consistent with an upper critical solution temperature (UCST) behavior. Holographic analyses of the same condensates revealed tight size and refractive index distributions of condensates at 30 and 42 °C. This was also different from DIC data, where we observed smaller condensates at 42 °C, potentially a surface attachment artifact. However, at 50 and 60 °C, we observed a skewed distribution containing a low refractive index population of large condensates and a high refractive index population of small condensates. The former population was not observed via DIC microscopy, potentially due to longer settling times due to low density. Specifically, the number density of the detected condensates dropped from 1.6 × 10^7^ mL^−1^ at 30 °C to 9.5 × 10^5^ mL^−1^ at 60 °C, which is consistent with the idea that PopZ condensates exhibit UCST behavior [3].

**Fig. 4.**
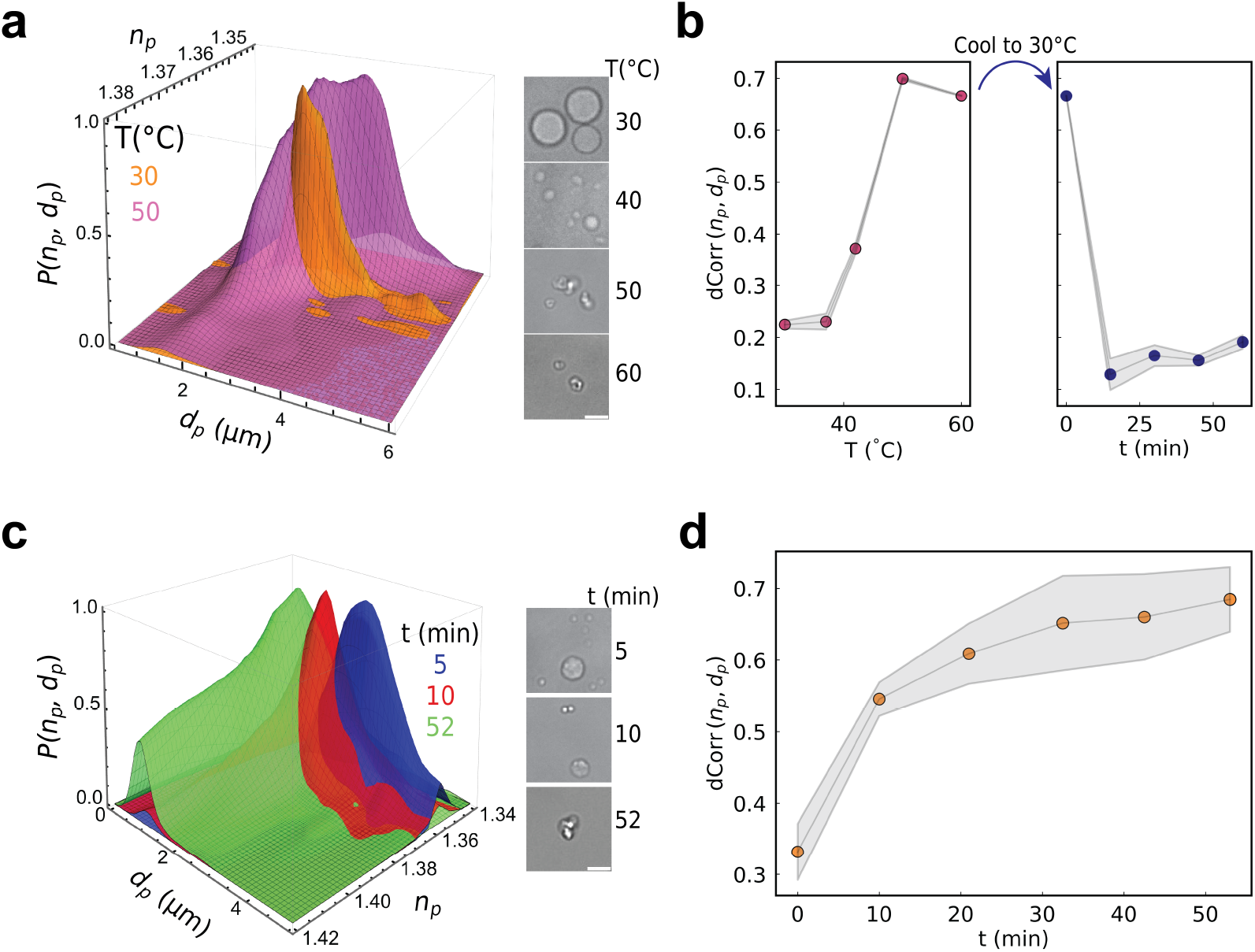
Out of equilibrium behavior of PopZ condensates. (a) Left panel: DIC microscopy images of PopZ condensates (5 µM at 150 mM Mg^2+^) at different temperatures (30 °C, 42 °C, 50 °C, and 60 °C), showing morphological changes with increasing temperature. Scale bar: 3 µm. Right panel: Surface plot of the refractive index (*n*_*p*_) against the diameter (*d*_*p*_) of the same condensates at two different temperatures (30 °C and 50 °C), with probability densities *P* (*n*_*p*_, *d*_*p*_) shown as the heights of the surfaces. At the two highest temperatures, the shapes of the *n*_*p*_ v *d*_*p*_ distributions are typical of condensates far from equilibrium. At lower temperatures, refractive indexes converge to a specific dense phase concentration value. (b) Distance correlation between refractive index and diameter dCorr(*n*_*p*_, *d*_*p*_) acts as a proxy for distance from equilibrium as a function of temperature, showing an increase at higher temperatures. Right panel: sharp reversal of the temperature-induced increase in dCorr(*n*_*p*_, *d*_*p*_) by returning the system to 30 °C, demonstrating a time-dependent recovery of dCorr(*n*_*p*_, *d*_*p*_) over 50 minutes. Shaded areas represent errors obtained by bootstrapping. (c) Left panel: DIC microscopy images of PopZ condensates as a function of time post lipoic acid addition. Scale bar: 3 µm. Right panel: Surface plot of the refractive index (*n*_*p*_) against the diameter (*d*_*p*_) of the same condensates. (d) Time-dependent behavior of the dCorr(*n*_*p*_, *d*_*p*_) as a function of time, before, and after lipoic acid addition, showing a sharp increase in dCorr(*n*_*p*_, *d*_*p*_) just after addition of lipoic acid followed by a gradual stabilization over time. Shaded areas represent error bars, computed by combining uncertainties from two sets of measurements.

We used distance correlation, as defined in Sec. 5, as a proxy for the distance of the system from equilibrium. Condensates far from equilibrium typically appear asymmetrical, as shown in the DIC micrographs in Fig. 4a. Such morphologies exhibit strong anti-correlations between *n*_*p*_ and *d*_*p*_, which are captured by measuring the distance correlation dCorr(*n*_*p*_, *d*_*p*_). Large values of dCorr(*n*_*p*_, *d*_*p*_) correspond to larger deviations from equilibrium. During the heating process dCorr(*n*_*p*_, *d*_*p*_) obtained from size distributions increased monotonically while upon cooling back to 30 °C, dCorr(*n*_*p*_, *d*_*p*_) converged to lower values as the condensates approached equilibrium (Fig. 4b). Next, we used lipoic acid, a compound previously shown to fluidize PopZ condensates [3] as a chemical perturbation. We monitored the condensate dissolution process as a function of time after adding 5 mM lipoic acid (Fig. 4c). In this case, we observed the distribution shift toward small-sized condensates with high refractive indexes.

This distribution could result from fluidization of large condensates, resulting in the formation of smaller, protein-rich assemblies that may be stabilized by hydrophobic interactions with lipoic acid. This interpretation is further strengthened by the observed increase in dCorr(*n*_*p*_, *d*_*p*_) immediately after the addition of lipoic acid, followed by a moderate increase as the system reaches its new equilibrium (Fig. 4d). Taken together, these data show the utility of holographic characterization, and provide a transferable analytical framework to study biomolecular condensates away from equilibrium.

### Excipient multivalency modulates PopZ condensate composition and dynamics

Building on the observation that polyvalent cations promote denser PopZ condensates and broaden the refractive index distribution (Fig. 2b), we next investigated the structural heterogeneity within individual droplets and its relationship to condensate dynamics. The hypothesis generated by holographic microscopy, that polyvalent cations introduce greater opportunities for structural heterogeneity was tested by examining both the spatial organization and temporal behavior of PopZ molecules within condensates. We applied single-molecule localization microscopy (SMLM) on condensates that were sparsely labeled with fluorescent PopZ [50]. By employing oblique illumination to further minimize perturbations, we observed individual fluorescent molecules within the condensates, which after localization and reconstruction, revealed distinct sub-diffraction clusters, as shown in Fig. 5a (top). The degree of clustering varied depending on the cation used, with more prominent clustering observed in Mg^2+^ and Pmm samples. We applied a density-based spatial clustering of applications with noise (DBSCAN) analysis to the single molecule data to assess the number of loalizations per cluster. DBSCAN analyses show that clusters inside condensates formed with Mg^2+^ and Pmm have a higher number of localizations compared to Spd^3+^ and Sp^4+^. In addition to nanoscale clusters, we detected a population of rapidly moving molecules, particularly in samples containing Spd^3+^ and Sp^4+^, which could not be precisely localized due to their fast motion in and out of the focal plane.

**Fig. 5.**
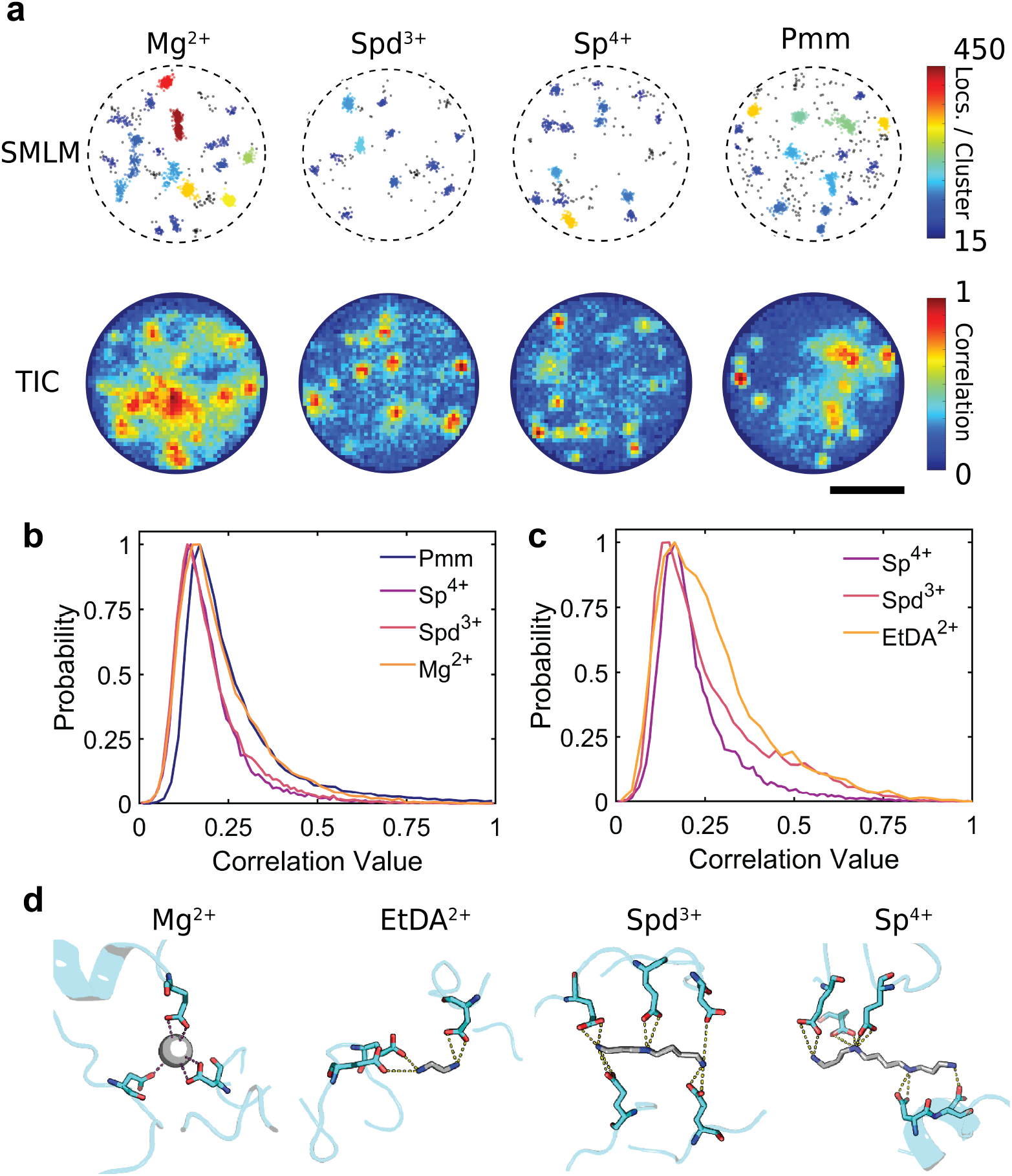
Effects of multivalent ions on PopZ condensate structure and dynamics. (a) Top: Single molecule localization microscopy and bottom: temporal image correlation analyses of PopZ condensates in the presence of various cations: Mg^2+^, Spd^3+^, Sp^4+^, and Pmm dendrimer. The condensates were labeled using 0.001 % (v/v) JF646 conjugated PopZ. SMLM highlights clustered localizations colored by the number of localizations in the cluster and outliers colored in black, while TIC maps regions of correlated molecular localizations within condensates. Scale bar: 2 µm. (b) Normalized frequency distributions of TIC correlation values for condensates formed with different multivalent ions. Comparison of Mg^2+^, Spd^3+^, SpSp^4+^, and Pmm reveals distinct shifts in molecular motion dynamics. (c) Comparison of EtDA^2+^, Spd^3+^, and Sp^4+^ at matched ionic strengths indicates decreased temporal correlation with higher valence of the multivalent cations. (d) Molecular interaction models showing representative binding configurations of PopZ with Mg^2+^ (gray sphere), EtDA^2+^, Sp^4+^, and Spd^3+^. Green dashed lines indicate hydrogen bonding and ionic interactions between PopZ and the multivalent cations.

Rapid motion of labeled PopZ molecules led to fluctuations in the fluorescent background within the condensates that were analyzed via temporal image correlation spectroscopy (TICS) [51]. Temporal correlation over a single time lag allowed us to differentiate dynamic from static regions, with blue regions in Fig. 5a (bottom) indicating higher molecular dynamics (low temporal correlation) and red regions representing static areas (high temporal correlation). Static regions within Spd^3+^ and Sp^4+^ correlated well with clusters of high density of molecules. In contrast, condensates formed using Mg^2+^ and Pmm showed some clusters that were dynamic despite having a high molecular density as measured via DBSCAN analyses. TICS analyses revealed that condensates formed with Sp^4+^ exhibited the highest molecular dynamics, followed by Spd^3+^ and Pmm, while Mg^2+^ displayed the lowest dynamics (Fig. 5b). However, this comparison was confounded by differences in ionic strength between the samples and the dendrimeric structure of Pmm, which makes direct interpretation challenging. To address this, we conducted a second experiment where the ionic strength was matched across all samples, and the valence of the multivalent cations was systematically varied, progressing from ethylenediamine dichloride (EtDA^2+^) to Spd^3+^ and Sp^4+^. In this experiment where the ionic strengths were controlled, EtDA^2+^ showed the lowest dynamics, closely followed by Spd^3+^ and Sp^4+^ (Fig. 5c). These results strongly suggest that the multivalency of the cationic excipients plays a key role in governing the dynamic interactions within PopZ condensates.

To further explore the molecular basis of these observations, we performed all-atom molecular dynamics (MD) simulations of PopZ in the presence of Mg^2+^, EtDA^2+^, Spd^3+^, and Sp^4+^. The simulations revealed distinct interaction profiles (Fig. 5d). Mg^2+^ exhibited a tightly coordinated network of electrostatic interactions with acidic residues within a small volume, while larger, multivalent molecules like Sp^4+^ showed spatially distributed hydrogen-bonding interactions between amino hydrogens and acidic residues, forming multiple binding sites across the molecule. Further, an analysis of the number of molecular interactions observed between each cation and PopZ chains shows that Mg^2+^ exhibits the highest interaction counts across all bond numbers, reflecting its strong and frequent interactions with protein residues (Fig. S5). In contrast, Sp^4+^, while having the lowest interaction counts for 1 and 2 bonds, dominates at higher bond numbers (5 to 6 bonds) due to its ability to form multiple simultaneous interactions. Among the multivalent ions, shorter chains like EtDA^2+^ show higher interaction counts at lower bond numbers, while longer chains such as Sp^4+^ and Spd^3+^ are better at higher bond numbers due to their extended structures and additional interaction sites. The distributed hydrogen bonding, lower frequency of molecular interactions, and lower interaction strength of hydrogen bonds compared to electrostatic interactions offered via Mg^2+^ likely explain the enhanced dynamics and compositional heterogeneity observed in condensates formed with higher valence excipients.

Our findings reveal that the valence of excipients used to trigger condensation is a critical determinant of structural heterogeneity and molecular dynamics in biomolecular condensates. By promoting distributed interactions within the condensate, higher valence cations create more opportunities for dynamic rearrangements. This insight into the role of multivalency in driving condensate behavior provides a deeper understanding of the physicochemical principles underlying condensate formation and regulation, with implications for material properties in biological and synthetic systems.

## Discussion

Measuring the precise size and composition of biomolecular condensates is essential to understand their formation, regulation, and impact on cellular processes. These measurements help differentiate functional condensates from pathological aggregates and assess bulk-like properties. Traditional methods rely on fluorescence labeling or surface attachment, which can alter condensate properties. Here, we report the use of holographic microscopy to measure the size and concentration of the dense phase of a model bacterial condensate, PopZ. This method is able to collect information for ∼ 5000 particles in ∼ 1 min, achieving greater precision and accuracy than conventional methods. Holograms of PopZ condensates revealed that dense phase concentrations are tunable through excipient and protein concentrations. Our results show that the initial protein concentration governs the resultant condensate sizes, recapitulating previous observations *in vivo* in a PopZ overexpression system [29]. The detection of bimodal size distributions in PopZ condensates highlights the unique ability of this measurement technique to resolve complex size distributions that other methods fail to detect, uncovering multiple growth mechanisms. While these mechanisms are not fully accounted for in the current model, the model nonetheless captures the essential aspects of condensate growth.

Time-dependent size distributions of PopZ condensates discerned via holography revealed growth dynamics different from previously reported exponential or powerlaw size distributions of condensates [14]. We applied a polyelectrolyte-attachment model [49] that predicts slow condensate growth, which was consistent with observations at late times. Using the distance correlation between index of refraction and size of condensate populations, we investigated the approach to equilibrium for PopZ condensates as well as their dissolution and reformation by temperature and lipoic acid. The data collectively illustrate the dynamic response of bacterial condensates to thermal and chemical stress, highlighting both immediate and reversible changes in their viscoelastic properties. Finally, our results demonstrate that excipient multivalency modulates the structural heterogeneity and molecular dynamics of PopZ condensates by enabling distributed interactions that drive dynamic rearrangements, highlighting the critical role of cation valence in condensate regulation. Importantly, this study illustrates the utility of holographic microscopy as a powerful hypothesis-generating tool, enabling non-invasive insights into condensate sub-structure that can be further tested and refined using complementary, minimally perturbative methods such as single-molecule localization and molecular modeling to uncover mechanisms of intra-condensate organization *in situ*.

Despite these advances, the method has limitations and relies on certain assumptions. The existing commercial implementation of holographic microscopy has a lower size limit of 0.5 µm, which means that condensates smaller than this limit are not detected (see Fig. 3a). When performed at a single wavelength, holographic microscopy provides a single value for the refractive index of each particle or droplet. This value must be interpreted with effective-medium theory [24, 25, 52] and modeling [53, 54] to infer the composition of a heterogeneous material such as a multi-component condensate. Future work will focus on studying two- and three-component condensates with the goal of estimating species-specific volume fractions to get relative occupancy of condensate scaffolds. Another area of research will be the application of holographic microscopy to study condensates that regulate enzymes, utilizing the dCorr(*n*_*p*_, *d*_*p*_) parameter to estimate the activity inside the condensate. Finally, comparison between conventional and holographic imaging modalities promises access to the relationship between condensate growth mechanisms and surface tension and therefore paves the way for a careful analysis of size distributions to infer interfacial energies that could be used for assessing condensate viscoelastic properties.

## Online methods

### Protein Expression and Purification

PopZ, tagged with a His_6_ epitope through a TEV protease cleavage site, was overexpressed in *E. coli* (BL21(DE3)) and purified under denaturing conditions [3, 30]. Bacterial cells were lysed in a buffer containing 8 M urea, 500 mM NaCl, 50 mM sodium phosphate (pH 7.0), 10 mM imidazole, and 200 mg mL^−1^ guanidinium chloride. His_6_-tagged PopZ was purified using Ni-NTA resin beads in a gravity column. The purified protein was refolded via dialysis and stored at −80 °C in a buffer containing 50 mM sodium phosphate (pH 7.0), 100 mM NaCl, and 10 % glycerol (v/v) until further use.

### Condensate Preparation

To prepare condensates, the His_6_ tag was removed from PopZ by incubating the purified protein with 0.03 M*/*M TEV protease. The sample was subsequently dialyzed using a 3.5 kDa MWCO membrane for 1 h at room temperature into a final buffer of 5 mM sodium phosphate (pH 7.0) and 10 mM NaCl.

Phase separation was induced by adding a multivalent cationic excipient—either MgCl_2_, spermidine trichloride, spermine tetrachloride, or PAMAM—to the protein solution. The mixture was incubated at 30 °C for 1 h, unless otherwise stated.

### Lipoic Acid Preparation and Addition

Lipoic acid solutions were prepared at 500 mM in 100 % DMSO and then diluted into 5 mM sodium phosphate (pH 7.0) and 10 mM NaCl to reach a final concentration of 50 mM. These solutions were further diluted into the condensate samples for a final lipoic acid concentration of 5 mM. During incubation, samples were mixed at 30 °C and 300 rpm on a standing shaker.

### Differential Interference Contrast (DIC) Microscopy Sample Preparation

To prevent surface interactions, all well plates used for DIC microscopy were pretreated with Tween-20. The wells were first flushed with N_2_ gas to remove dust and debris, then coated with 1 % Tween-20 and incubated at 42 °C for 1 h. After incubation, the wells were washed five times with water at twice the volume of the Tween-20 solution. Finally, the wells were dried with N_2_ gas and used within the same day.

### Condensate Imaging with DIC

Differential Interference Contrast (DIC) microscopy was used to analyze phase boundaries and condensate size distributions. For phase boundary analysis, condensate samples were added to a Tween-20 treated well, sealed with an adhesive, and imaged using a multi-well plate imager (Cytation 5) equipped with a 60x air objective preheated to 30 °C. The use of a plate reader enabled rapid phase space imaging and phase boundary determination. Images were acquired approximately 15 min after adding the samples to the well plate.

For size distribution analysis, condensate samples were added to a Tween-20 treated well plate and imaged using an inverted microscope (Nikon Ti2) equipped with DIC optics, an oil immersion objective (Nikon PlanApo, 100×, 1.45 NA), and a sCMOS camera (Photometrics Prime 95B) with a system magnification of 0.11 µm*/*pixel. After 2 min of incubation, ten 1200 pixel × 1200 pixel Z-stacks were acquired with a 40 ms exposure time per frame for each sample within a 3 min time window.

For size distribution analysis, an optimal Z-height was determined for each region of interest (ROI) where condensates appeared with high contrast. Image segmentation was performed in MATLAB to estimate droplet diameters, while ensuring that out-of-focus condensates were excluded. Errors in diameter estimation were primarily due to variations in the optimal Z-height for each droplet and uncertainty in droplet edge detection thresholds. Systematic offsets from potential substrate adhesion effects were not included in the uncertainty analysis.

### Condensate Imaging and Characterization with Holographic Microscopy

Holographic imaging was performed using a commercial instrument (Spheryx xSight) [18]. 30 µL aliquots of condensate samples were transferred into a commercial microfluidic channel (Spheryx xCell8) with a height of 50 µm and a width of 500 µm. A pressure-driven Poiseuille flow transported the condensate particles through the channel at a maximum speed of (3 ± 1) mm s^−1^. Dispersed particles entrained in the flow were illuminated by a 450 nm laser, and the resulting holograms were recorded.

The instrument analyzed each hologram using a generative model [18, 20] based on the Lorenz-Mie theory of light scattering [32], extracting both the particle diameter, *d*_*p*_, and refractive index, *n*_*p*_. A single measurement on a micrometer-scale spherical particle yielded diameter estimates with a precision of ±2 nm and refractive index measurements accurate to within ±1 × 10^−3^ [18, 20, 22]. At particle concentrations around 10^7^ particles*/*mL, statistical sampling of thousands of particles could be achieved and analyzed in under 15 min. The pipeline for holographic imaging and automated analysis is depicted schematically in Fig. 1b.

### Dye Labeling and Confocal Imaging of Condensates

PopZ-TEV-His_6_ was labeled using either BODIPY-NHS or JF646-SE dyes. PopZ-His_6_ and the specific NHS-dye were incubated together in 5 mM sodium phosphate (pH 7.0) and 10 mM sodium chloride. The reaction mixture was kept in the dark with slow mixing for 6 h, followed by dialysis into 50 mM sodium phosphate (pH 7.0), 100 mM sodium chloride, and 10 % glycerol (v/v). Protein concentration was determined using the Pierce™ Bradford Plus Reagent, while dye concentration was determined from the absorbance spectrum of the solution. This conjugation protocol yielded labeled protein samples with a degree of labeling of 0.8 to 1 dye molecule per protein. The dye-labeled protein was flash-frozen and stored at 80 °C.

For dye-labeled protein condensate reconstitution, labeled PopZ was mixed with unlabeled protein at specific volume percentages. Samples were prepared as described in the section for DIC imaging.

To measure relative fluorescence intensities of condensates, samples were added to a Tween-20 treated well plate and imaged after 5 min of incubation to allow for settling. Imaging was performed using a confocal laser scanning microscope (Abberior), and at least three 80 µm × 80 µm ROIs were collected per condition. Images were processed using bespoke Python programs. Thresholding was applied to isolate condensates, and the mean fluorescence intensity inside each condensate (*F*_in_) was recorded. To determine background fluorescence, pixels inside the condensates were set to NaN, and the mean intensity of the remaining image was computed as *F*_out_. The fluorescence ratio, *F*_in_*/F*_out_, was calculated for each condition, as shown in Fig. S3.

To determine relaxation times, 5 % (v/v) JF646-labeled PopZ was added to unlabeled PopZ in a 5 mM sodium phosphate (pH 7.0) and 10 mM sodium chloride buffer to achieve a final protein concentration of 5 µM. Phase separation was induced with 150 mM magnesium chloride. After a 1 h incubation at 30 °C, condensates were added to a Tween-20 treated well plate and imaged immediately using the Aberrior STEDY-CON microscope in the *x*-*z* plane. Five image sequences of 100 frames over 200 s were collected for ROIs measuring 70 µm to 90 µm (x-axis) by 10 µm to 14 µm (z-axis).

### Single Molecule Localization Microscopy of PopZ Condensates

PopZ labeled with JF646 by NHS-ester conjugation as previously described was added to an unlabeled sample of PopZ at 0.001 % v/v. After condensate formation, samples were added to a Tween-20 treated 384-well glass-bottom plates (number 1.5, Sigma). The condensates fixed on the surface of the well were imaged within 10 minutes after addition to the well plate. 1000 frames were collected at an exposure time of 20 ms for each ROI. At least six 256 × 256 pixel ROIs were collected for each sample. The number of condensates analyzed in each case were: Mg^2+^ (23), EtDA^2+^ (10), Spd^3+^ (16), Sp^4+^ (20), and Pmm (31).

ThunderSTORM [55] analysis with normalized Gaussian visualization from FIJI was performed on representative condensates for each sample, as shown in Fig. 5. For temporal image correlation spectroscopy (TICS), condensate images were first thresholded to include intensity values from pixels only within the condensate using a threshold *T* calculated by the following:

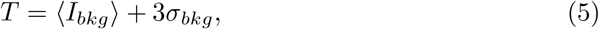

where the mean intensity ⟨*I*_*bkg*_⟩ and standard deviation *σ*_*bkg*_ of the background were used. Intensities at each pixel of a video were normalized to *z* scores using the following equation,

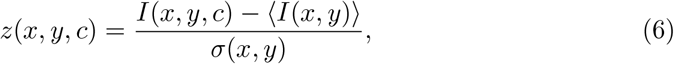

where *z*(*x, y, c*) represents the *z* score at a specific frame number *c* and pixel *x, y. I*(*x, y, c*) represents the raw intensity value at the frame number c and ⟨*I*(*x, y*) ⟩ is the average intensity at that pixel. *σ*(*x, y*) is the standard deviation of intensity values at that pixel. These *z* scores were then processed further to determine the average correlation value at that pixel with a specified time lag *τ* with units in frames:

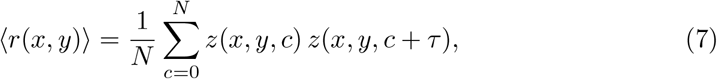

where *N* = 1000 denotes the total number of frames collected for each video. This process was performed for all pixels within a condensate, and the average correlation values per pixel were plotted either as a heat map or enveloped histogram. To account for photobleaching, a correction factor was calculated based on previous calculations using TICS [56]. Briefly, average intensities of the condensates of a single field of view were plotted as a function of time *t*. The intensities were normalized and fit to an exponential decay curve,

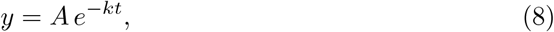

where *y* represents the normalized average intensity and *k* is the bleaching decay constant. Average correlation values were then multiplied by the correction factor,

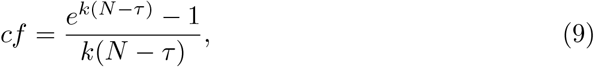

to produce the final correlation values, which were normalized and spatially resolved as in Fig. 5.

### All-atom Molecular Dynamics Simulations of PopZ and Multivalent Cations

All-atom MD simulations were performed to capture details of molecular interactions between PopZ and multivalent ions.The simulation system was built by randomly placing 14 chains of PopZ protein in a 20 nm cubic box, which was subsequently solvated by TIP4P water [57]. Keeping ionic strength of 2 M, multivalent ions, including Mg^2+^, EtDA^2+^, Spd^3+^, and Sp^4+^, were introduced into the box to mimic the experimental condition, while chloride ions were added to neutralize the system. The Mg^2+^ box contained 242 324 water molecules, 3279 Mg^2+^ ions, and 6152 Cl^−^ ions. The EtDA^2+^ box included 229 256 water molecules, 3279 EtDA^2+^ molecules, and 6152 Cl^−^ ions. The Spd^3+^ box was composed of 230 125 water molecules, 1640 Spd^3+^ molecules, and 4514 Cl^−^ ions. Finally, the Sp^4+^ box consisted of 234 521 water molecules, 984 Sp^4+^ molecules, and 3530 Cl^−^ ions. Periodic boundary conditions were applied in all three dimensions for all systems. Energy minimization was performed using the steepest descent algorithm followed by 100-ps equilibration, with a step size of 2 fs, under NVT conditions at 300 K. Protein position restraints were applied during NVT run. Further NPT equilibration was performed for 100 ps with a 2-fs time step, using the Parrinello-Rahman barostat for pressure coupling to maintain a pressure of 1 bar, without any restraints on protein position. Two production MD runs were carried out for 10 ns and 83.4 ns, with a time step of 2 fs. Energy and compressed coordinate data were saved every 10 ps (5000 steps) for subsequent analysis. All simulations were conducted using GROMACS 2023.3 [58] and the AMBER99SB force field [59].

Hydrogen bonds between negative residues on PopZ chains and multivalent cations were identified using a cutoff distance of 3.9 Å between acceptors and donors. Ionic bonds between negative residues on PopZ chains and Mg^2+^ ions were defined using a cutoff distance of 5 Å [60]. Each frame of the trajectory file was analyzed, looping over all multivalent ions to identify interactions and quantify the number of protein interactions per ion. The total number of frames where a certain number of interactions, 1 to 6, was counted and presented in a logarithmic-scale plot with the number of interactions on the *x*-axis and the log of total number of interaction counts on the *y*-axis. By analyzing interaction modes and the total number of interaction counts, we predict the effect of multivalent ions on the stability of the protein system. Notably, the short and long runs exhibited similar trends, reinforcing the consistency of the observed dynamics.

## Supplementary information

Methods for protein concentration measurement, correlation metrics and supplementary figures (S1-S5) are provided in the supplement.

## Acknowledgments

The authors thank members of the Grier and Saurabh research groups, and Prof. Alexander Grosberg (NYU) for helpful discussions and inputs. This study was supported by National Institutes of Health through award 1R35GM157103-01 to SS, and by the National Science Foundation (NSF) through award DMR-2104837 to DGG. SW was supported by an NSF funded REU site in chemical biology at NYU Chemistry. The xSight instrument used for this study was acquired as shared instrumentation with support from the MRSEC program of the NSF under award DMR-1420073.

## Declarations

DGG is a founder of Spheryx, Inc., the company that manufactures xSight for Total Holographic Characterization, including the instrument used for the present study.

## Supplementary Information

### Holographic Measurement of Protein Concentration in the Condensed Phase

Total Holographic Characterization (THC) (the commercial implementation of holographic microscopy with microfluidics) yields a value for the refractive index, *n*_*p*_, of each condensate droplet with part-per-thousand precision [18, 25]. This information can be used to infer precise values for the volume fraction and absolute concentration of proteins in the dense phase of a phase-separated solution. This method has not been reported previously and represents a new application area for holographic microscopy.

The measured refractive index is related to the volume fraction, *ϕ*, of protein in the dense phase through Maxwell Garnett effective-medium theory [24, 25, 61],

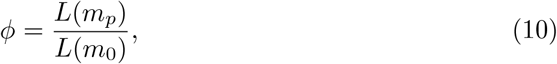

where *m*_*p*_ = *n*_*p*_*/n*_*m*_ is droplet’s refractive index relative to that of the medium, *m*_0_ = *n*_0_*/n*_*m*_ is the relative refractive index of the protein itself, and

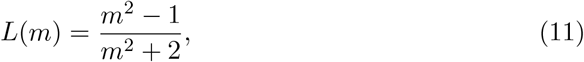

is the Lorentz-Lorenz function. The volume fraction of protein within the condensate is proportional to the protein concentration, *c*,

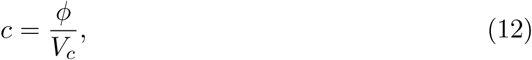

where *V*_*c*_ is the volume of a single protein chain. The refractive index of the medium, *n*_*m*_, is measured conveniently with an Abbe refractometer (Abbe-3L Refractometer, Fisher Scientific). Measuring the dense-phase protein concentration also requires precise values for the intrinsic refractive index of the protein, *n*_0_, and the associated single-chain volume, *V*_*c*_.

The intrinsic refractive index of the protein can be obtained from its amino acid sequence using effective-medium theory together with tabulated data [62] for the refractive indexes, 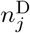, and specific volumes, *v*_*j*_ of the pure amino acids. Tabulated values are reported for the a vacuum wavelength of 589 nm, which corresponds to the sodium D line. The result for an *N* -amino-acid protein,

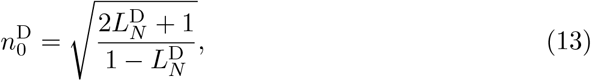

is a function of the volume-weighted Lorentz-Lorenz factor,

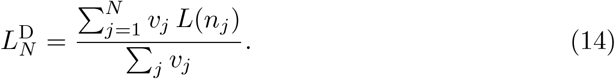

This result is scaled for use at other wavelengths, *λ*, using a standard result for protein refractive-index increments [63],

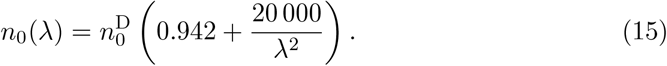

This approach yields *n*_0_ = 1.689 ± 0.001 for the intrinsic refractive index of PopZ at the imaging wavelength used for holographic characterization, *λ* = 450 nm. This value can be used in Eq. (10) to convert measurements of droplets’ refractive indexes into estimates for the volume fraction of protein within the droplets, as presented in Fig. 2a. The typical single-droplet uncertainty in the estimated volume fraction is Δ*ϕ* = ±0.2 %, based on propagation of uncertainties in 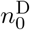 and the parameters in Eq. (15) [63].

The largest uncertainty in estimating protein concentrations can be ascribed to the value used for the single-chain volume, *V*_*c*_. This is the volume associated with the protein’s intrinsic light-scattering properties, which is substantially smaller than the volume subtended by the protein’s tertiary structure. We obtain an estimate for *V*_*c*_ by using an Abbe refractometer to measure the refractive index, *n*_*s*_, of bulk protein solutions as a function of protein concentration, *c*. These values are converted into estimates for the volume fraction of protein in the bulk using the computed value of 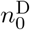 and the measured refractive index of the buffer, *n*_*m*_, as inputs to Eq. (10) as inputs. Typical results for PopZ are plotted as discrete points in Fig. S3a. The dependence of *n*_*s*_ on *c* yields an estimate for the optical volume, *V*_*c*_, through Eq. (12). The result for PopZ, *V*_*c*_ = (260±67) nm^3^, agrees with independent estimates obtained numerically using the CALVADOS model [64], which also are plotted in Fig. S3a. We use the calibrated value of *V*_*c*_ to convert holographically measured values of the droplet refractive index into the estimates for the dense-phase PopZ concentration that are reported in Fig. 2. The same protocol can be used to measure the densephase concentration in other protein solutions undergoing phase separation, and can be generalized to measure macromolecular concentrations in heterogeneous condensates.

### Jensen-Shannon Divergence

To quantify the reproducibility of size measurements using holographic characterization and DIC, we compute the Jensen-Shannon Divergence (JSD) scores between sets of measurements from the same method. Size distributions between two runs using the same technique provides an estimate for the reproducibility of the method. The probability distributions are most similar when the JSD approaches 0. The JSD itself is derived from the Kullback Leibler Divergence (KLD), which, for probability distributions *p*_1_ and *p*_2_ is given as

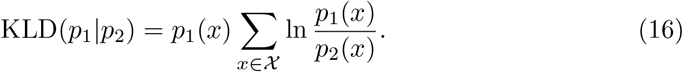

If *m* is a mixture distribution,

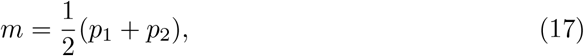

the JSD is computed as [35]

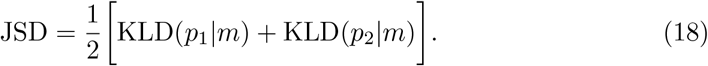

We computed the JSD scores between holographic microscopy runs and obtained consistently lower scores across the range of Mg^2+^ when compared to the JSD scores between DIC runs. As a benchmark, we compared a probability distribution obtained from holography with a uniform distribution with the same range [*a, b*], where *a* and *b* are the minimum and maximum sizes respectively measured using holographic microscopy.

### Distance Correlation

Distance correlation scores between condensates’ refractive indexes and sizes were used as a proxy for distance from equilibrium. The distance correlation score is defined as [65]

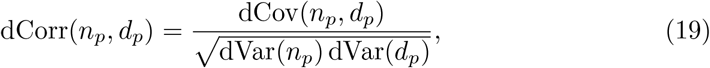

the distance covariance between *n*_*p*_ and *d*_*p*_, normalized by the square root of the product of distance variances. The distance covariance between two quantities *X* and *Y* is defined as

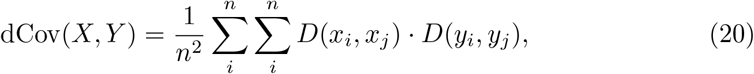

where *D*(*x*_*i*_, *x*_*j*_) is the double-centered distance matrix for quantity *X* of size *n*, and the distance variance is the covariance with itself

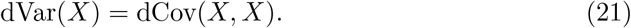

Unlike the more commonly used Pearson correlation, the distance correlation does not assume a linear relationship between quantities. Additionally, the distance correlation score dCorr(*n*_*p*_, *d*_*p*_) is constrained between 0 and 1, where dCorr(*n*_*p*_, *d*_*p*_) = 0 strictly applies only to independent variables.

## Supplementary Figures

**Fig. S1.**
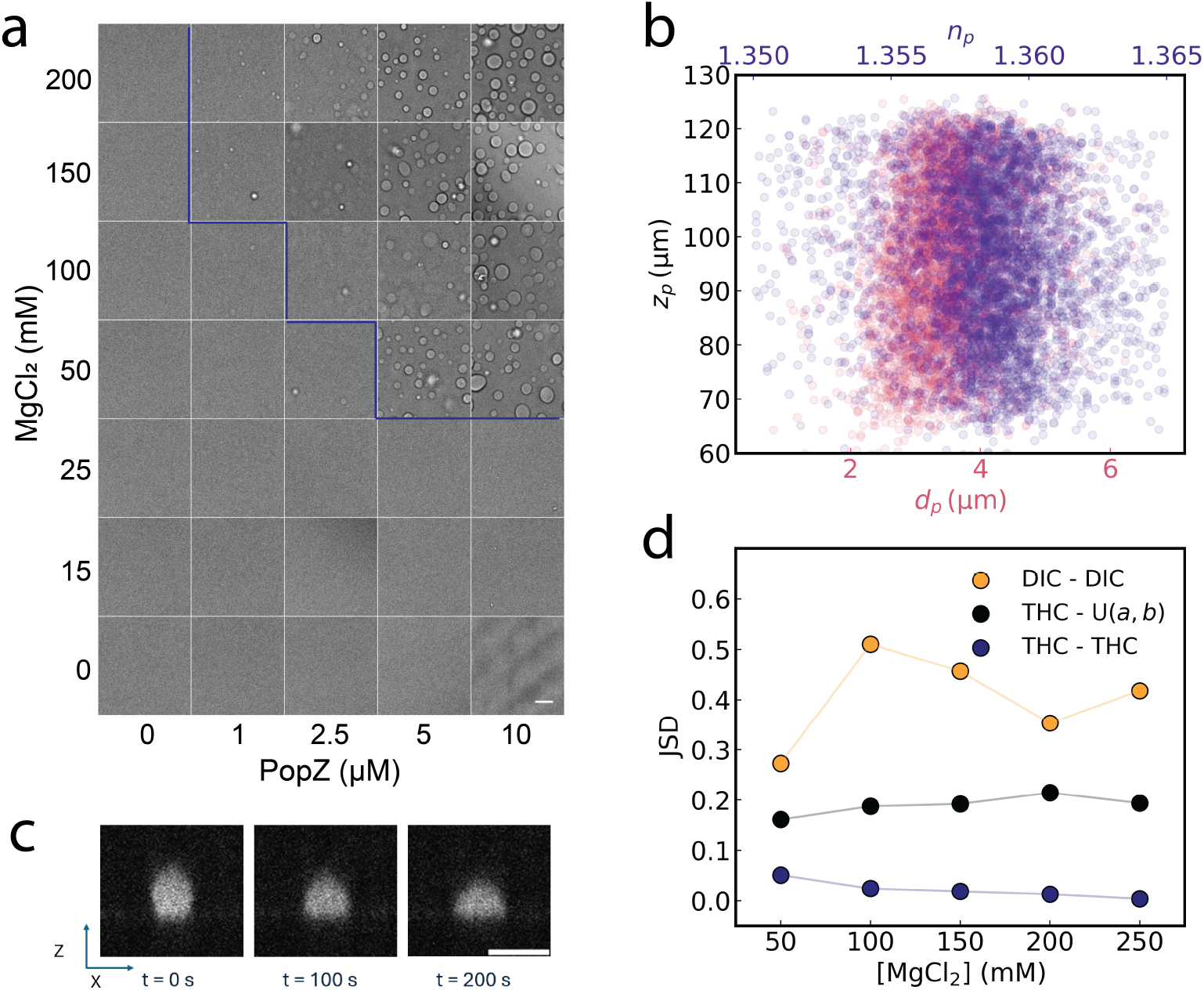
Characterization of PopZ condensates droplets and their phase behavior. (a) Phase behavior of PopZ as a function of protein and Mg^2+^ concentration measured via DIC. Scale bar: 5 µm. (b) Scatter plot demonstrating that the measured diameter (*d*_*p*_) and refractive index (*n*_*p*_) of individual PopZ condensates is independent of their axial position (*z*_*p*_). (c) Time-lapse images showing the dynamics of a single PopZ condensate (labeled using 5 % Bodipy-PopZ) over 200 s, visualized in the *x*-*z* plane using confocal microscopy. Scale bar: 5 µm. (d) The Jensen-Shannon Divergence (JSD) is computed for the size measurements shown in panel A. Comparing JSD scores between holography measurements and DIC measurements reveals that holography measurements are more reproducible than DIC measurements. The JSD score between a size distribution obtained from holography and one that follows a uniform distribution with the same mean and standard deviation is shown for reference.

**Fig. S2.**
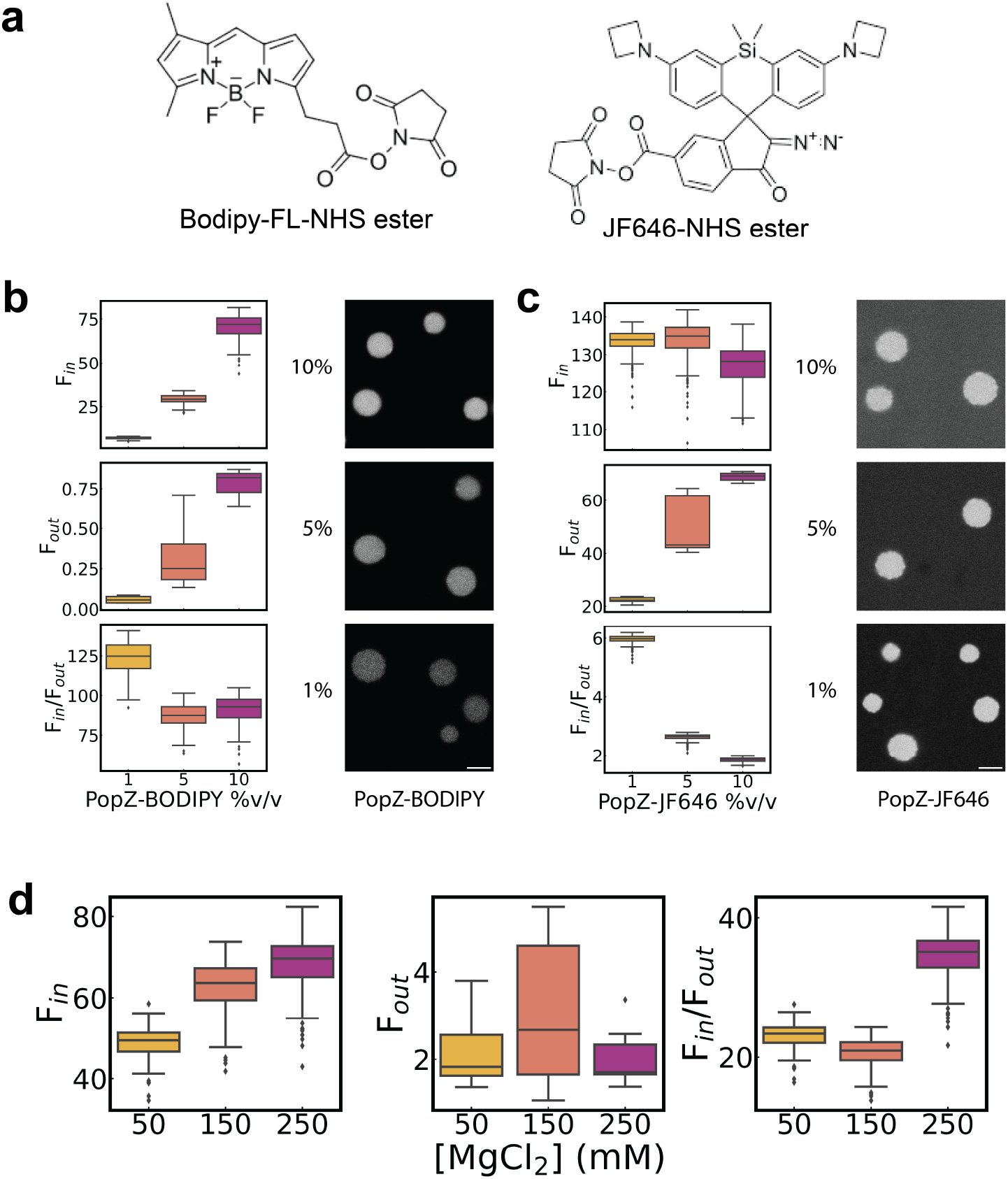
Characterization of PopZ Condensates via fluorescence microscopy. (a) Structure of the NHS-ester derivatives of JF646 and Bodipy-FL, the dyes used in this study. (b) Left panels: Boxplots of fluorescence intensity in (*F*_*in*_) and outside (*F*_out_) condensates at different percent v/v additions of dye-labeled PopZ (left; PopZ-BODIPY-FL to an unlabeled PopZ sample. 5 µM PopZ and 200 mM Mg^2+^ were prepared for each experiment. Relative concentrations of dense phase were determined using the ratio of *F*_in_*/F*_out_. Representative confocal images are shown. (c) Same data as in (b) but with PopZ condensates labeled with PopZ-JF646. Scale bar: 5 µm. (d) Boxplots of *F*_in_ and *F*_*in*_ of PopZ-BODIPY labeled condensates (5% v/v) as a function of Mg^2+^ concentration. Relative concentrations of dense phase PopZ were determined using the ratio of *F*_in_*/F*_out_.

**Fig. S3.**
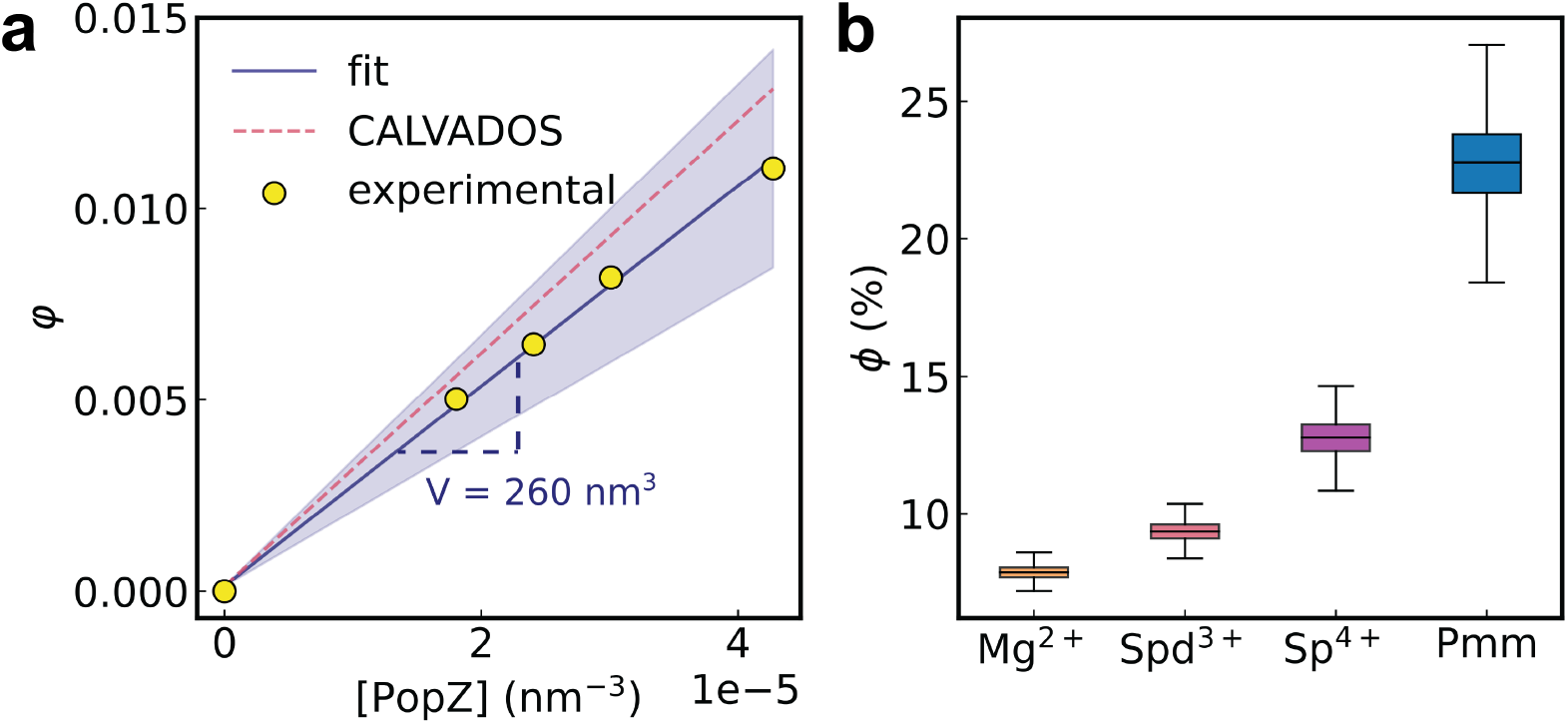
Concentration and volume fraction measurements of PopZ condensates. (a) Plot of calculated volume fractions *ϕ* as a function of the volume of a single PopZ chain determined experimentally by measuring the index of refraction of PopZ monomer solutions of known concentrations using an Abbe refractometer. The blue shaded area shows the uncertainty in our estimation of the volume. These experimental results are consistent with estimates from CALVADOS. (b) Volume fractions of condensates formed using different excipient multivalent ions.

**Fig. S4.**
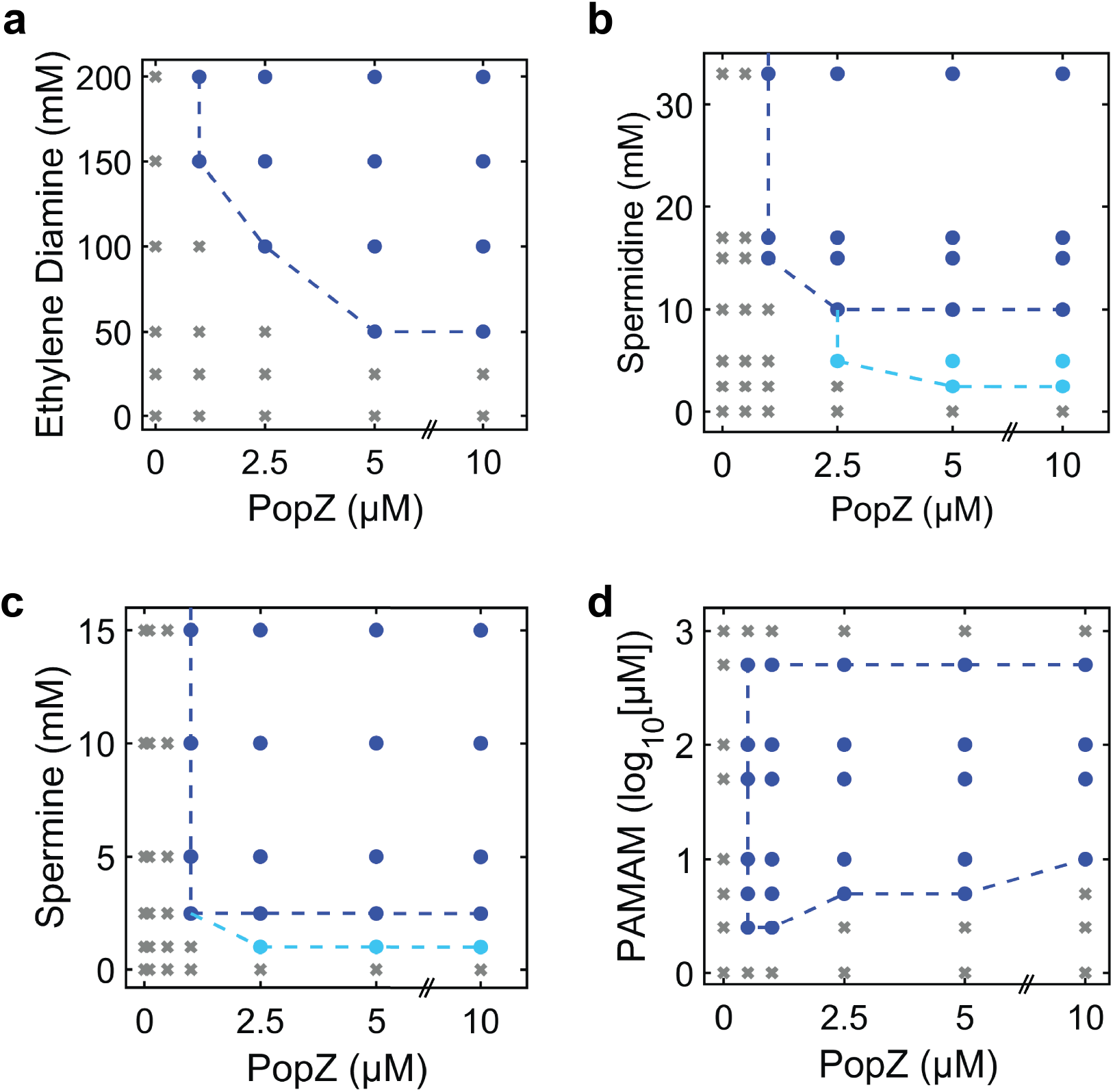
Phase diagrams of PopZ in the presence of polyamines. (a) Ethylene diamine (EtDA^2+^), (b) Spermine (Sp^4+^) (c) Spermidine (Spd^3+^), and (d) PAMAM dendrimer (Pmm). Phase diagrams were measured using a multiwell imager, with PopZ reconstituted in 5 mM Sodium Phosphate (pH 7.0) and 10 mM NaCl. Data were collected using DIC microscopy. Gray X-marks indicate that no phase separation was observed. Spherical condensates were observed in conditions labeled with dark circles. Light blue circles indicate aspherical structures. Boundaries are drawn for reader’s convenience.

**Fig. S5.**
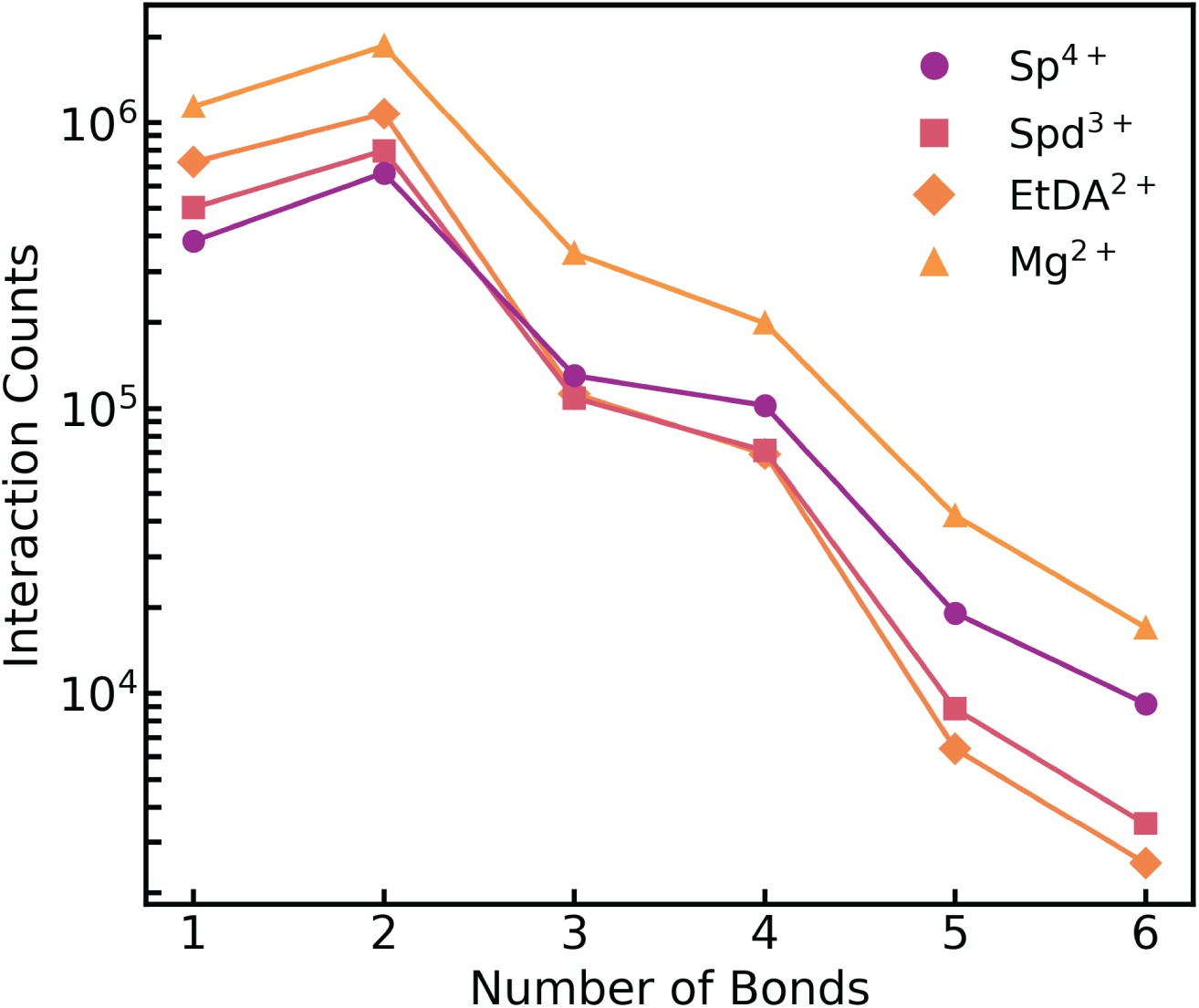
Molecular interactions of PopZ with various excipients observed via MD simulation. Total number of molecular interaction counts observed for each excipient ion with PopZ chain, for different number of bonds (ionic or hydrogen). These data are from the analysis of simulations run for a total time of 83.4 ns. Lines connecting the symbols are for reader’s convenience.

